# *In Silico* Driven Prediction of MAPK14 Off-Targets Reveals Unrelated Proteins with High Accuracy

**DOI:** 10.1101/2020.07.24.219071

**Authors:** Florian Kaiser, Maximilian G. Plach, Christoph Leberecht, Thomas Schubert, V. Joachim Haupt

## Abstract

During the discovery and development of new drugs, candidates with undesired and potentially harmful side-effects can arise at all stages, which poses significant scientific and economic risks. Most of such phenotypic side-effects can be attributed to binding of the drug candidate to unintended proteins, so-called off-targets. The early identification of potential off-targets is therefore of utmost importance to mitigate any downstream risks. We showcase how the combination of knowledge-based *in silico* off-target screening and state-of-the-art biophysics can be applied to rapidly identify off-targets for a MAPK14 inhibitor. Out of 13 predicted off-targets, six proteins were confirmed to interact with the inhibitor *in vitro*, which translates to an exceptional hit rate of 46%. For two proteins, affinities in the lower micromolar range were obtained: The kinase IRE1 and the Hematopoietic Prostaglandin D Synthase, which is entirely unrelated to MAPK14 and is involved in different cell-regulatory processes. The whole off-target identification/validation pipeline can be completed as fast as within two months, excluding delivery times of proteins. These results emphasize how computational off-target screening in combination with MicroScale Thermophoresis can effectively reduce downstream development risks in a very competitive time frame and at low cost.

## Introduction

In response to extracellular stimuli, Mitogen-activated protein kinase (MAPK) pathways control a multitude of cellular processes from cell growth [1] to DNA repair [2]. These stimuli trigger the release of mitogen peptides that bind to cell surface receptors. This binding signal is then relayed to the internal machinery of the cell via a signal cascade and eventually activates a variety of effector kinases. MAPKs themselves are serine/threonine kinases and are activated in the MAPK pathway via different pathways [3]. One of them is the p38 pathway that is responsive to stress stimuli such as heat, osmotic pressure, and radiation [4]. MAPK14 (also called p38-*α*) is ubiquitous in many cell types and instrumental for DNA repair [5, 6]. In recent years it has become clear that MAPK14 is also involved in autophagy [7] and is thus a viable target for cancer therapy [8, 9].

### Motivation

Identifying kinase inhibitors is generally not too complicated; many kinase targets are often bound by inhibitors with nanomolar affinity. The major challenge during the development of kinase inhibitors is to reach the necessary high selectivity and specificity to avoid undesired, harmful, and potentially lethal side effects that arise from binding to other proteins. In this study, we showcase how the combination of knowledge-based algorithms with efficient biophysical testing allows for the rapid identification of such undesired off-targets with high accuracy. We selected the kinase inhibitor SB203580, which was originally designed for MAPK14. SB203580 has already been linked to the inhibition of several other proteins, e.g., inhibition of RAF1 *in vitro* [10], phosphoinositide-dependent protein kinase 1 [11], cardiac stress-activated protein kinases [12], and bromodomain containing protein [13]. At the time of writing, the ChEMBL database [14] lists 498 distinct proteins with measured activities against SB203580, emphasizing the critical role of off-targets for this inhibitor. However, this comprehensive list is not yet complete.

### Computational Off-Target Prediction

The computational prediction of off-targets can follow different routes. Chemical similarity screening allows to infer off-target effects by searching for known compounds that are chemically similar to the drug candidate [15, 16], which can identify off-targets for which no protein structure data is known. However, such approaches are often imprecise and predictions can be substantially improved, if protein structure data is available. Considering metabolic pathway information is another promising tool [17], but fails predicting arbitrary drug-target combinations if there are no paths in the underlying pathway network that connect drug and target [18]. Recent advances in computational off-target prediction compose meta-approaches, aggregating knowledge from different sources, mainly via text-mining of literature and structure activity relationship (SAR) data [19]. Methods that are based on artificial intelligence depend on the availability of high-quality training data. While there are many experimental measurements for positive drug-target interactions, negative samples often fall short [18]. Knowledge-based methods, such as the PharmAI *DiscoveryEngine,* can mitigate these limitations.

### Off-Target Identification Service

In a joint effort, we developed a computational workflow to predict off-targets in *silico* and validate these predictions in *vitro* with unmatched speed (see Figure 1). The underlying PharmAI *DiscoveryEngine* exploits hidden similarities between binding sites of otherwise unrelated (i.e. non-homologous) proteins [20] and similarities in the non-covalent interaction patterns of drug-target complexes [21]. This unique combination of features and the application of artificial intelligence allow for the prediction of proteins that potentially bind to a given compound with unmatched speed and thus cost efficiency. Recently, we have applied the same technology to discover novel cGMP-dependent 3’,5’-cyclic phosphodiesterase (PDE2) inhibitors with an outstanding hit rate of 6% [22]. The starting point for a PharmAI Off-Target screening is either the protein structure of a desired on-target, the chemical structure of a small molecule, or a combination thereof. In these cases, the method can identify potential off-targets rapidly. The prediction covers proteins where structure data is available from the Protein Data Bank (PDB) [23] or external resources. With this approach we currently cover approximately 20,000 human proteins and 95% of all known human drug targets.

**Figure 1.**
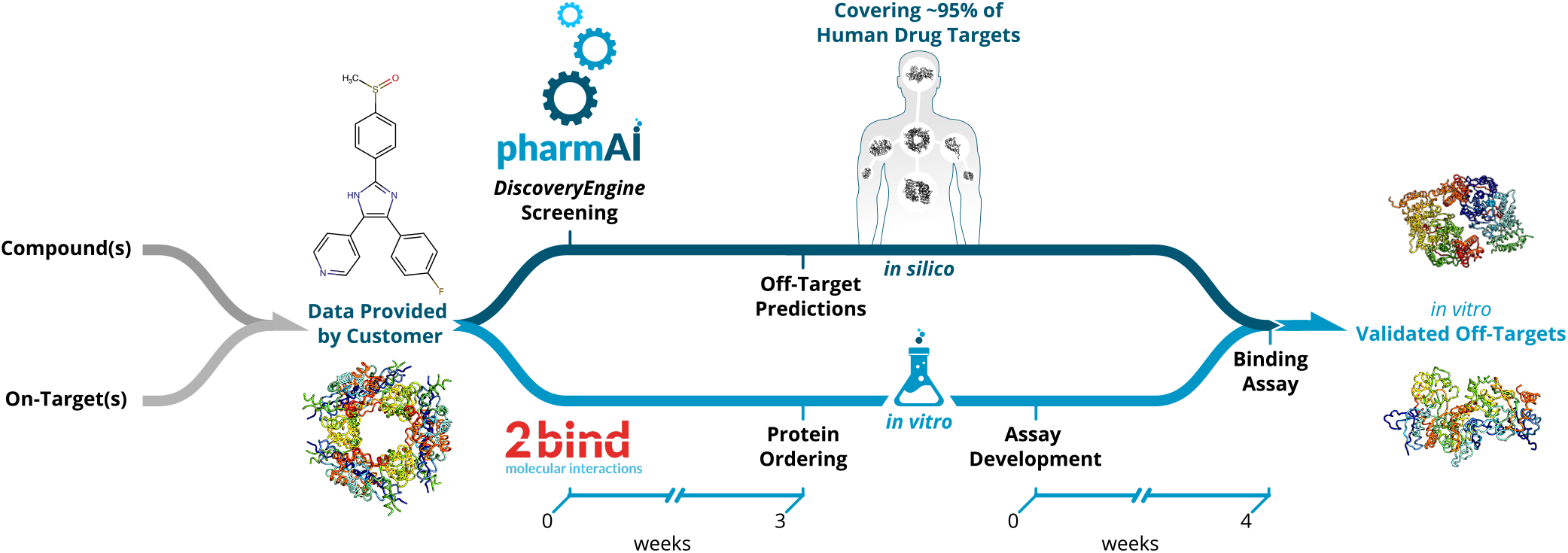
Workflow of the PharmAIo2bind *Off-Target Identification Service.* Upon the definition of the compound(s) and/or on-targets(s) of interest, *in silico* and *in vitro* pipelines are launched in parallel. The PharmAI *DiscoveryEngine* rapidly predicts potential off-targets within one week, covering 95% of the known human drug targets. Predicted proteins are then purchased and assay development starts as soon as the *in silico* data are ready.

### MicroScale Thermophoresis (MST)

Successful application of the PharmAI·2bind *Off-Target Identification Service* requires fast, efficient, robust, and accurate *in vitro* validation of the computational predictions. Moreover, validation should consume as little protein as possible, because potentially many predicted off-targets have to be tested and usually there are no resources and time for lengthy production of large protein quantities. A method that combines speed and efficiency with ultra-low sample consumption and that can deliver the binding affinity (*KD*) of the compound of interest to the potential off-targets is MST. The 2bind MST service is fully compatible with lab automation (e.g. LabCyte Echo acoustics, contact-less compound preparation) to achieve quick results. For example, for an off-target analysis only 6 *μ*l of the compound of interest and only 50 *μ*g of each predicted off-target protein are required. MST relies on a unique measurement principle (Figure 2a): The predicted off-target proteins are labeled with a special fluorescent dye. A general property of fluorescence is that its intensity decreases with increasing temperature. This effect is called TRIC. Now this fluorescence change can be further manipulated by the binding of a ligand to the labeled target molecule (binding in close proximity to the dye or via conformational changes of the target). Together with an additional readout principle, thermophoresis, the MST signal is sensitively affected by various different molecular parameters, which usually all change upon binding of a ligand to a target molecule. The output of MST are classical and well-described ligand-binding dose-response curves, which can be fitted to yield the steady-state affinity (*K_D_*) of an interaction (Figure 2b).

**Figure 2.**
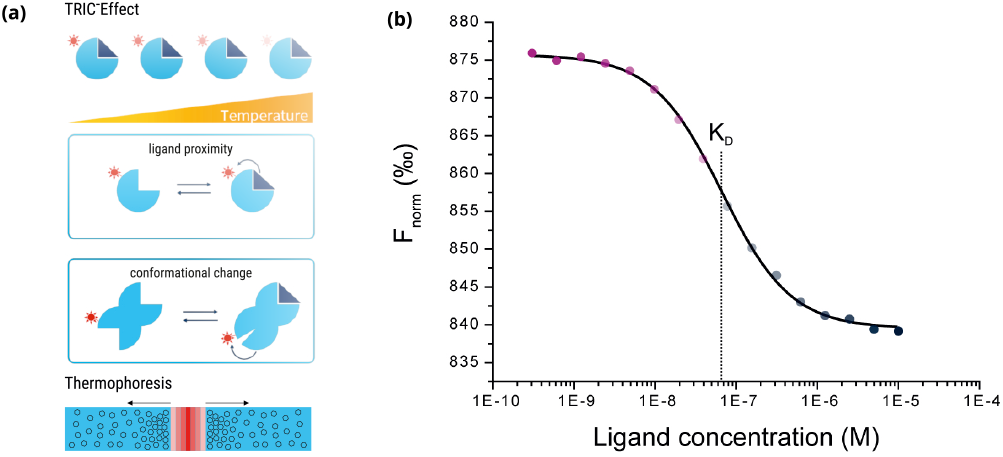
Principle and output of MicroScale Thermophoresis (MST). (**a**) Schematic illustration of the temperature-related intensity change (TRIC) effect (top and middle) and the thermophoretic movement of molecules in a temperature gradient (bottom). (**b**) The calculated F_norm_ values from the MST traces are dose-dependent and can be well described by the law of mass action. A fit of these values returns the dissociation constant *K_D_* of the interaction.

## Results and Discussion

Starting from SB203580 co-crystallized with MAPK14, we applied the PharmAI *DiscoveryEngine* and identified 167 proteins as potential off-targets for SB203580. Out of these proteins, 56 were already known to bind SB203580 *in vitro.* Importantly, bromodomain containing protein was among the reported off-targets. This non-kinase protein is already known to bind SB203580 [13] which emphasizes the power of the *DiscoveryEngine* to pick-up proteins that are functionally and evolutionary not related to the actual target. From the predictions that have not been previously linked to SB203580 (according to activity measurements in the ChEMBL database [14]), the top 25 predictions were short-listed for *in vitro* validation. Of these 25,13 proteins were commercially available in sufficient quality and were purchased in small quantities (usually <100 *μ*g) for biophysical testing. SB203580 showed binding to six of these proteins in the 2bind *Off-Target* MST assay, translating to an exceptional hit rate of 46% (Table 1). The validated targets included the kinase IRE1 with a *K_D_* of 2.05 *μ*M to SB203580 and the enzyme HPGDS, which bound SB203580 with a *K_D_* of 30.87 *μ*M. Figure 3 shows the MST curves for both off-targets together with docking poses of SB203580. The MST response amplitudes were sufficiently high and the signal/noise level was very good for both tested proteins. Thus, the *K_D_* values could be determined from the data fit with high accuracy and certainty. The other four validated off-targets were weak binders with *K_D_* values >500 *μ*M (Figure S1). For these proteins, higher concentrations of SB203580 would have to be tested to resolve a full dose-response curve. All of the positively validated off-targets share no sequence similarity with a sequence identity of <10% to MAPK14 according to a pairwise alignment [33].

**Figure 3.**
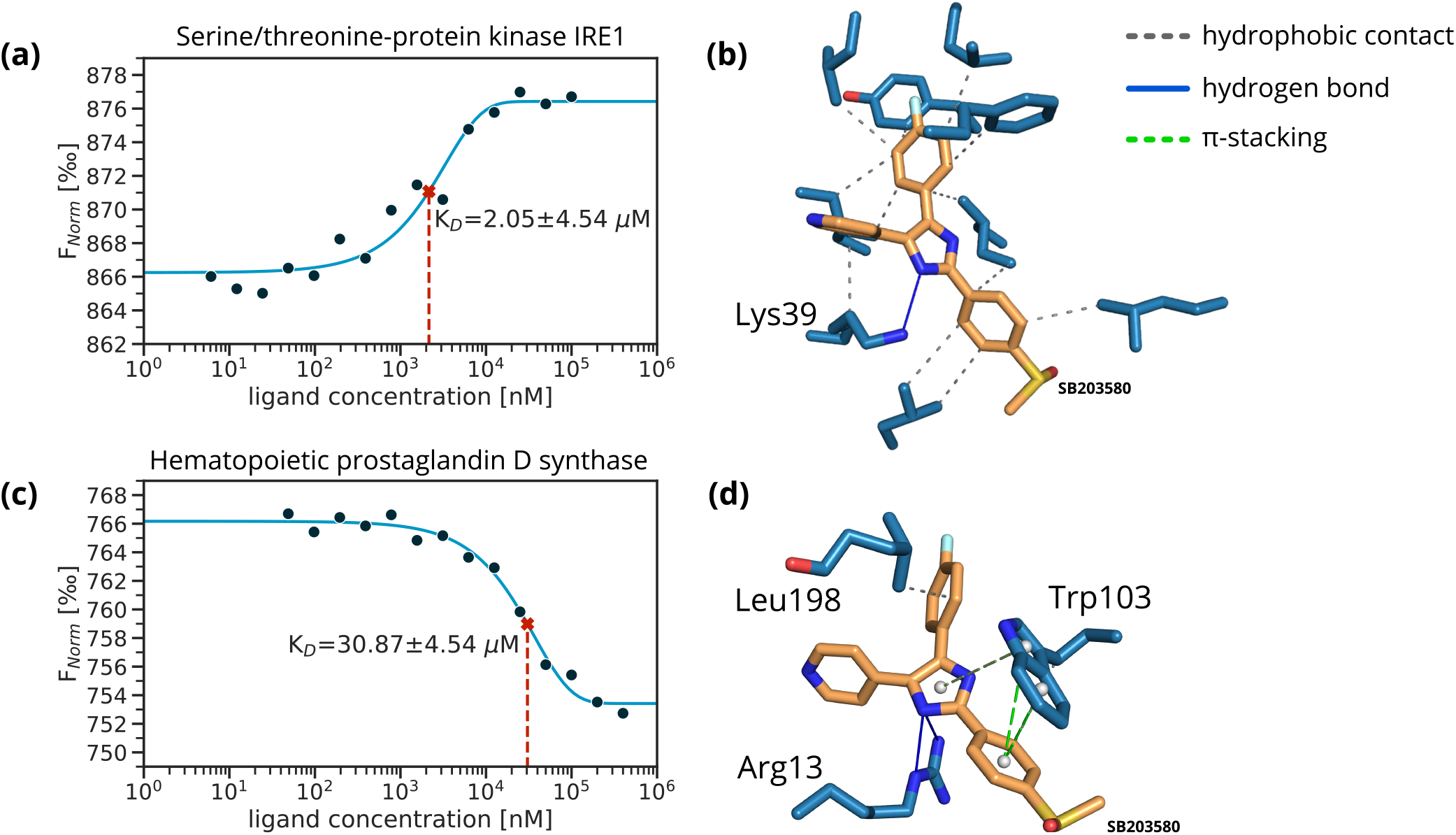
MST dose-response graphs and docking poses for Serine/threonine-protein kinase/endoribonuclease IRE1 (IRE1) and Hematopoietic prostaglandin D synthase (HPGDS). (**a**) Dose-response graph for the kinase IRE1. (**b**) The best docking pose of SB203580 obtained with AutoDock Smina and a flexible docking protocol [30, 31] for IRE1 (PDB-ID: 4U6R, −11.40 kcalomol^−^1). Annotated non-covalent interactions are shown [32]. (**c**) Dose-response graph for HPGDS. (**d**) The best docking pose of SB203580 obtained with AutoDock Smina and a flexible docking protocol [30, 31] for HPGDS (PDB-ID: 2VCW, −11.50 kcalomol^−^1). Annotated non-covalent interactions are shown [32].

**Table 1.**
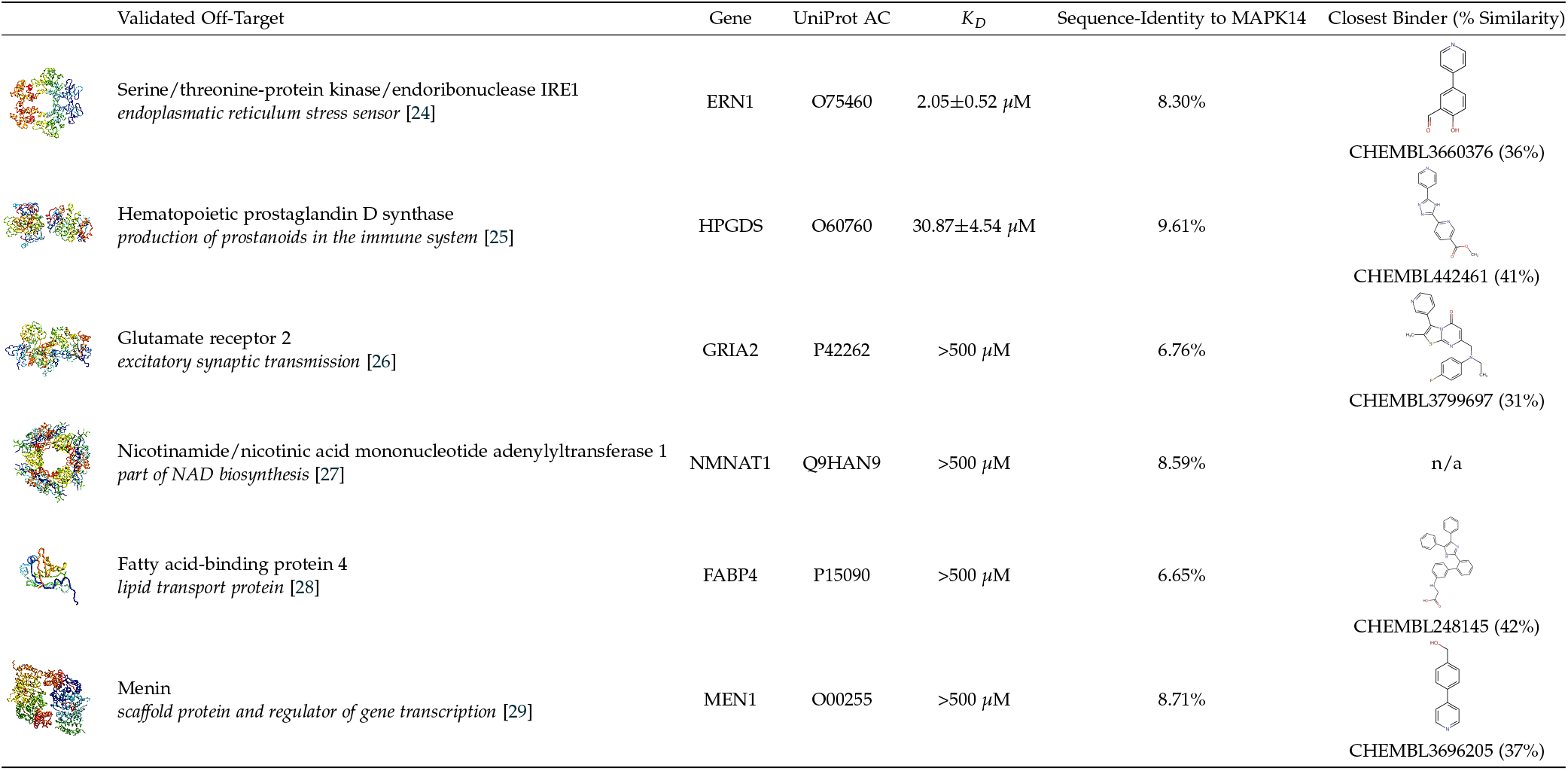
Validated off-targets for the kinase inhibitor SB203580. For each protein, the determined *K_D_* to SB203580 and the sequence identity to MAPK14 is given. Additionally, the chemically most similar compound to SB203580 with reported activity [14] for each protein is shown.

| Validated Off-Target | Gene | UniProt AC | $K_D$ | Sequence-Identity to MAPK14 | Closest Binder (% Similarity) |
| --- | --- | --- | --- | --- | --- |
| Serine/threonine-protein kinase/endoribonuclease IRE1<br>endoplasmatic reticulum stress sensor [24] | ERN1 | O75460 | $2.05 \pm 0.52 \mu\text{M}$ | 8.30% | CHEMBL3660376 (36%) |
| Hematopoietic prostaglandin D synthase<br>production of prostanoids in the immune system [25] | HPGDS | O60760 | $30.87 \pm 4.54 \mu\text{M}$ | 9.61% | CHEMBL442461 (41%) |
| Glutamate receptor 2<br>excitatory synaptic transmission [26] | GRIA2 | P42262 | $>500 \mu\text{M}$ | 6.76% | CHEMBL3799697 (31%) |
| Nicotinamide/nicotinic acid mononucleotide adenylyltransferase 1<br>part of NAD biosynthesis [27] | NMNAT1 | Q9HAN9 | $>500 \mu\text{M}$ | 8.59% | n/a |
| Fatty acid-binding protein 4<br>lipid transport protein [28] | FABP4 | P15090 | $>500 \mu\text{M}$ | 6.65% | CHEMBL248145 (42%) |
| Menin<br>scaffold protein and regulator of gene transcription [29] | MEN1 | O00255 | $>500 \mu\text{M}$ | 8.71% | CHEMBL3696205 (37%) |

We investigated whether the identified off-targets could have been found by a naive chemical similarity search of SB203580. For each of the off-targets we searched for the most similar known compound to SB203580 – defined by chemical fingerprints [34] – with reported activity in the ChEMBL database [14]. All of the known binders have chemical similarities of <50% to SB203580. Furthermore, a list of all molecules in ChEMBL was created ranked by the chemical similarity to SB203580. For each of these compounds, associated targets with measured activities were retrieved. None of the newly identified and validated off-targets appeared within the first 1000 ranks. Conclusively, these proteins could thus not have been found by a chemical similarity screening of SB203580 against the ChEMBL database without testing at least 1000 proteins *in vitro.*

In general, the correct and fast identification of off-targets is of high importance. On average, drugs have about 11 off-targets with IC_50_ values below 10 *μ*M [35]. Off-target prediction is therefore a highly valuable tool to learn about potential side effects of drugs, as well as opportunities for drug repositioning. In a large scale study Lounkine *et al.* [36] observed that about 10% of their tested compounds bound to nearly half of their targets. These results indicate that many off-targets remain undiscovered across various stages of the drug development pipeline, leading to eventual failure of approval and thus a huge financial loss.

## Conclusions

We showcased how the synergy between highly accurate knowledge-based *in silico* off-target prediction with the PharmAI *DiscoveryEngine* and fast and ultra-low sample consumption biophysical testing with the 2bind *Off-Target* MST assay can efficiently identify and validate off-targets at large scale. During the early development of new drugs, a tremendous reduction of downstream risks and costs can be achieved with this service. Compounds predicted to bind to many targets can be classified as highly promiscuous and therefore potentially dismissed. Knowing the target classes can shed light on whether the compound will be a specific binder. Furthermore, the comprehensive knowledge of potential binding partners besides the primary target can guide and aid molecular design by comparing their binding pockets. The presented methodology allows to fine-tune target selectivity right from the very beginning of the drug development pipeline.

## Methods

### Chemical Similarity

Chemical similarities were computed using Extended-Connectivity Fingerprints [34] with highly optimized parameters. ChEMBL version 26 was used to identify existing inhibitors.

### Computational Off-Target Detection

The PharmAI *DiscoveryEngine* analyzed the binding site of MAPK14 regarding non-covalent interaction patterns as well as geometrical and physicochemical properties in nine PDB structures (PDB-IDs: 1A9U, 1PME, 2EWA, 3GCP, 3MPA, 3OBG, 3UIB, 3ZS5, 5AR4), which are co-crystallized with SB203580. In a next step, these properties were converted into profiles describing the binding characteristics of SB203580. Subsequently, the derived profiles were screened against PharmAI’s data warehouse of protein and small molecule data. As a result, a hit list of 167 proteins was obtained, out of which 56 proteins were already known to bind SB203580 according to the ChEMBL database [14]. From the remaining 111 proteins, 13 were selected (out of the top 25) for biophysical validation with MST based on protein availability and quality.

### *In vitro* validation of off-target predictions

The 13 selected, potential off-targets were sourced from commercial vendors. The six *in* vitro-validated off-targets were: IRE1, Hoelzel Diagnostika (11905-H20B); HPGDS, bio-techne (6487-PS-010); GRIA2, bio-techne (9526-UR); NMNAT1, Antibodies Online (ABIN2745771); FABP4, Antibodies Online (ABIN1096606); Menin, Antibodies Online (ABIN4989089). All proteins were controlled for quality by nanoDSF thermal unfolding profiling using a Tycho NT.6 instrument (NanoTemper Technologies, Munich, Germany). SB203580 was purchased from Selleckchem (Cat. S1076). Proteins were labeled with a 647 nm, red-fluorescent MST dye (NHS coupling chemistry). 16-concentration serial dilutions of SB203580 were prepared via contactless, high-precision, acoustic dilution (LabCyte Echo platform). The MST analysis was performed on an NT.115pico instrument (NanoTemper Technologies, Munich, Germany) with a “pico-red” detector setting and “medium” MST-power setting. MST data were exported with MO.AffinityAnalysis (v2.3, NanoTemper Technologies, Munich, Germany) using a one-second MST on-time interval set between 5 and 10 seconds and otherwise standard settings. Dose-response curves were fitted to a 1:1 *K_D_* model derived from the law of mass action.

## Funding

Parts of the results presented in this study were obtained during a feasibility study funded by the Sächsiche Aufbaubank (SAB) with funds from the European Regional Development Fund.

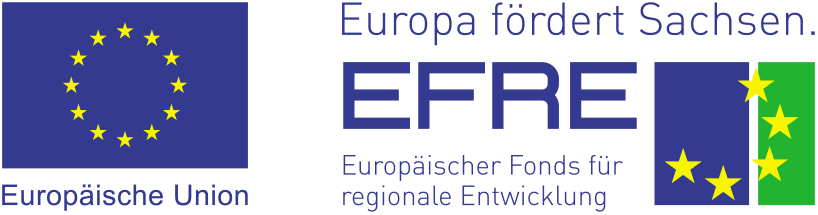

## Supporting Information

**Figure S1.**
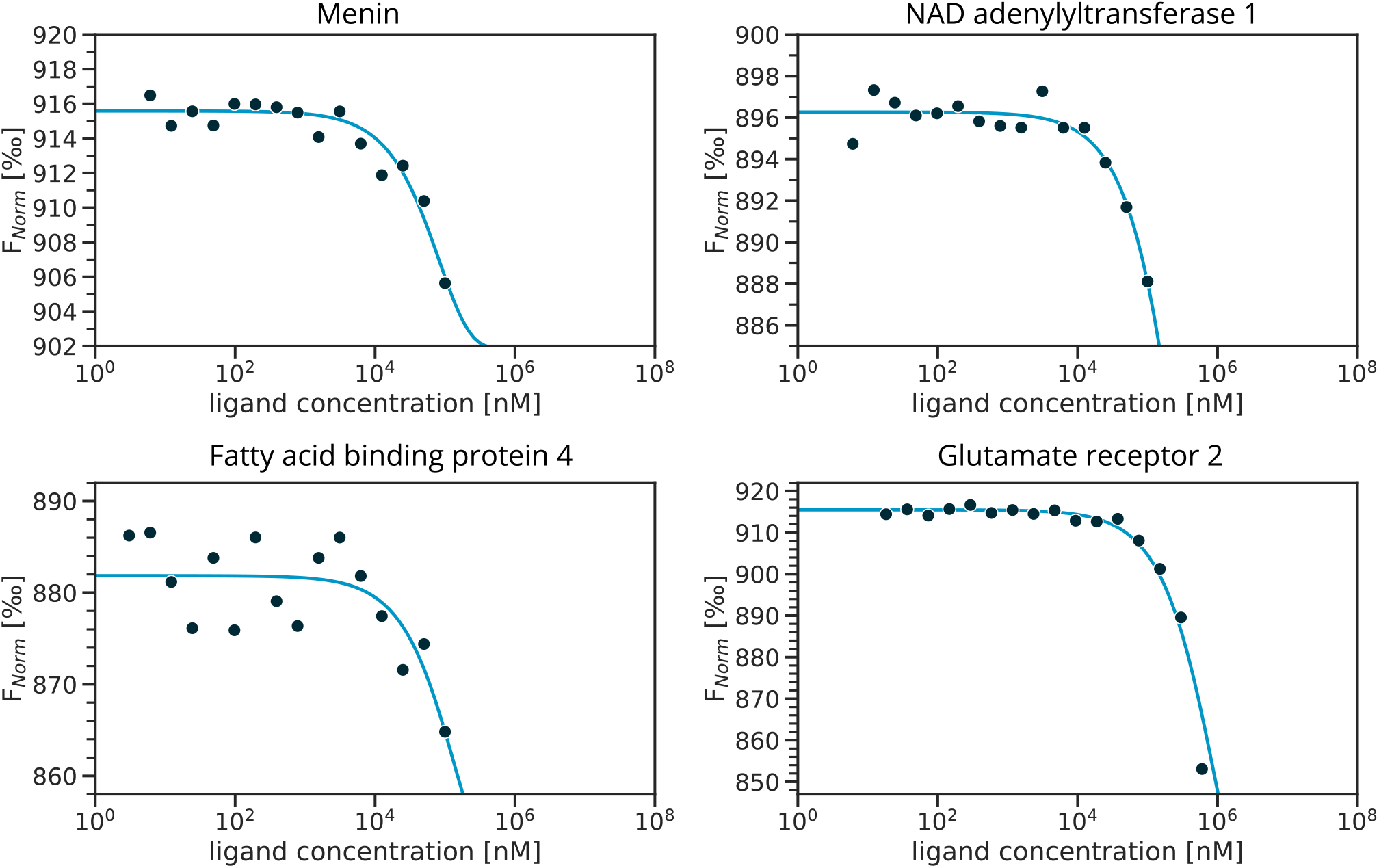
MST dose-response graphs for off-targets with weak binding to SB203580. The data indicate potential weak binding with *K_D_* values >500 *μ*M; a full curve-fit was not possible, because no bound plateau was reached at the maximum tested ligand concentration of 1 mM.

## Notes

### Competing Interest Statement

The authors have declared no competing interest.

